# Zika and dengue but not chikungunya are associated with Guillain-Barré syndrome in Mexico: a case-control study

**DOI:** 10.1101/2020.01.08.898403

**Authors:** Israel Grijalva, Concepción Grajales-Muñiz, César González-Bonilla, Victor Hugo Borja-Aburto, Martín Paredes-Cruz, José Guerrero-Cantera, Joaquín González-Ibarra, Alfonso Vallejos-Parás, Teresita Rojas-Mendoza, Clara Esperanza Santacruz-Tinoco, Porfirio Hernández-Bautista, Lumumba Arriaga-Nieto, María Garza-Sagástegui, Ignacio Vargas-Ramos, Ana Sepúlveda-Núñez, Omar Israel Campos-Villarreal, Roberto Corrales-Pérez, Mallela Azuara-Castillo, Elsa Sierra-González, José Alfonso Meza-Medina, Bernardo Martínez-Miguel, Gabriela López-Becerril, Jessica Ramos-Orozco, Tomás Muñoz-Guerrero, María Soledad Gutiérrez-Lozano, Arlette Areli Cervantes-Ocampo

**Author notes:** Corresponding author; (IG).

## Abstract

**Background:** Zika, dengue and chikungunya viruses (ZIKV, CHIKV and DENV) are temporally associated with neurological diseases such as Guillain-Barré syndrome (GBS). Because these three arboviruses coexist in Mexico, the frequency and severity of GBS could theoretically increase. This study aimed to determine the association between these arboviruses and GBS in a Mexican population, and to establish the clinical features of the patients, including the severity of the infection.

**Methodology/Principal findings:** A case-control study was conducted (2016/07/01-2018/06/30) in hospitals from the Instituto Mexicano del Seguro Social. Serum and urine samples were collected to determine the exposure to ZIKV, DENV and CHIKV by RT-qPCR and serology (IgM). The association between arboviral infection and GBS was analysed with χ^2^ or exact Fisher tests and Kruskall-Wallis folloed by Mann Whitney U test for comparing symptomatology. A p ≤ 0.05 was considered as significant. Ninety-seven GBS cases and 184 controls were included. Evidence of association of GBS with ZIKV acute infection (OR, 8.04; 95% CI, 0.89–73.01, p =0.047), as well as ZIKV recent infection (OR, 16.68; 95% CI, 2.05–135.31; p = 0.001) and Flavivirus (ZIKV and DENV) recent infection (OR, 5.8; 95% CI, 1.8-18.8; p = 0.002) was observed. Cases with GBS associated with ZIKV demonstrated a greater impairment of functional status, a higher percentage of mechanical ventilation and mortality. Dizziness, ataxia and low blood pressure showed statistically signifant differences in cases with GBS associated with DENV. Co-infection cases were not observed.

**Conclusions/Significance:** According to the laboratory results, an association between ZIKV or ZIKV and DENV infection in patients with GBS was found. Cases of GBS associated with ZIKV but not with DENV exhibited a more severe clinical picture. Cases with co-infection were not found.

**Author summary:** Dengue (DENV), chikungunya (CHIKV), and Zika (ZIKV) are considered as emerging or re-emerging viruses. In recent years, these viruses have produced great epidemics in tropical climate urban centers, and have been associated with neurological manifestations, including Guillain-Barre syndrome (GBS), which causes muscle weakness, unstable gait, and decreased or absent musculoskeletal reflexes. This study aimed to investigate the association between these viral infections and GBS. A case and control study was conducted nationwide, including 97 cases of GBS and 184 controls matched by age, gender, and locality, but without the disease. This study showed the positive association between GBS cases and ZIKV or ZIKV and DENV infection. GBS cases associated with ZIKV showed a more severe clinical picture (more impairment of functional status and incapacity, a higher percentage of mechanical ventilation, and mortality). GBS associated with DENV cases seemed to show more dizziness, ataxia, and low blood pressure. Finally, the symptoms of ZIKV or DENV suspected disease before the development of GBS were similar to some previous reports. The impact of the interaction of these three arboviruses, particularly ZIKV, on the health of the Mexican population was less than expected. The Mexican experience could be useful for other populations.

## Introduction

Arboviruses are a significant cause of human disease worldwide [1]. Viruses such as Zika (ZIKV), dengue (DENV) and chikungunya (CHIKV) produce large epidemics in tropical climate urban centres [1] and have expanded recently with outbreaks of considerable scale in the Western Hemisphere [2].

Several neurological manifestations, including Guillain-Barré syndrome (GBS), are associated with ZIKV, DENV or CHIKV [3-5]. Moreover, these arboviruses have been found to produce co-infection related to neurological diseases in some patients [6,7].

GBS is an acute immune-mediated inflammatory polyradiculoneuropathy that typically produces predominantly distal symmetrical muscular weakness, unsteady gait, and hyporeflexia or areflexia [8]. About two-thirds of patients with GBS have reported a history of *Campylobacter jejuni*, cytomegalovirus, Epstein-Barr virus or *Mycoplasma pneumoniae*, as well as influenza A virus and *Haemophilus influenzae* infections. Recently, an association with ZIKV, CHIKV and DENV has been reported as well [3-5,8,9]. Mortality remains around 8% [10]. In Mexico, a mortality rate of 5-10% was documented in 2014 [11].

It has been known that these three arboviruses have coexisted in Mexico since 2015. On the one hand, the pre-existing immunity to DENV could strengthen ZIKV infection (both flaviviruses) by increasing the severity of the disease or co-infections [6,7,12,13]. On the other hand, given their coexistence, there is a probability of predominance of any of the three arboviruses depending on their interactions, especially since the Mexican population had not been exposed to ZIKV earlier. Considering these facts, the aim of this study was to identify the association of GBS with these arboviruses in a Mexican population, the demographic and clinical characteristics of the patients, the severity of the infection and co-infection.

## Methods

### Study design and participants

A case-control study was conducted to identify the role of ZIKV, DENV and CHIKV infection in the development of GBS in Mexico, from July 1, 2016, to June 30, 2018. The protocol was authorised by the National Committee of Scientific Research of the Instituto Mexicano del Seguro Social (IMSS). GBS patients and control subjects were selected from IMSS hospital units nationwide. The study was based on the Institutional Epidemiological Surveillance of Communicable Diseases system. The participation of the Researches was decided voluntarily.

Patients developing GBS were identified prospectively by a second level physician due to flaccid paralysis. If necessary, patients were sent to a third level care hospital. Patients were evaluated by a neurologist, who diagnosed GBS based on the clinical features and neurological manifestations of the disease, as well as cerebrospinal fluid (CSF) analysis and electrodiagnostic test results following international criteria [14]. Medical records were reviewed and diagnostic tests, including cerebrospinal fluid, neuroimaging and electrophysiological studies, when available, were conducted to establish the clinical characteristics of the disease and confirm the diagnosis of GBS. Patients who met levels 1-3 of diagnostic certainty according to the Brighton Collaboration criteria were considered as eligible and were invited to participate in the protocol.

The control group was formed by two individuals for each GBS-case recruited from the same hospital where cases were evaluated, including both healthy individuals and patients with a non-febrile disease (no fever documented 48 hours before enrolment). Controls were chosen as a non-probabilistic sample of individuals paired by age (± 5 years), gender and place of residence with the corresponding case, and enrolled within seven days from the date of the GBS case diagnosis. Clinical and demographic data were collected directly from the patients or their relatives. The information of the patient included age, gender, place of residence, clinical history of comorbidities, signs and symptoms, duration and severity of the disease with GBS Disability Score and Medical Research Council Scale for Muscle Strength (MRC) and treatment. GBS cases and controls were interviewed for more information about risk factors and exposure during the last two months before the onset of the neurological symptoms.

The neurophysiological evaluation included motor and sensory nerve conduction studies of upper and lower extremities peripheral nerves. A lumbar puncture was performed by standard technique to obtain CSF for cytological and cytochemical analysis.

After the interviews, serum and urine samples were collected to determine the exposure to ZIKV, DENV and CHIKV during the first seven days after the onset of the neurological picture. If the individuals agreed to participate in the study, blood and urine samples were collected from each control as well.

The following definitions were established to determine the type of ZIKV, DENV or CHIKV infection:

Acute infection by ZIKV, DENV or CHIKV was defined as a positive result to the reverse transcriptase real-time polymerase chain reaction (RT-qPCR) in either serum or urine samples.

Evidence of recent infection by ZIKV, DENV or CHIKV was defined as a positive RT-qPCR result in serum or urine samples, or an anti-ZIKV, anti-DENV or anti-CHIKV immunoglobulin M (IgM) detected in serum.

### Laboratory analysis

Diagnostic tests were performed at the Central Laboratory of Epidemiology, La Raza National Medical Center, IMSS, Mexico City, following the epidemiological surveillance guidelines and in compliance with the national regulations established by the Mexican Ministry of Health.

Viral RNA extraction from serum and urine samples was carried out using the QIAmp Viral RNA Mini Kit (Cat. 52906, Qiagen, Hilden, Germany) or the MagNA Pure LC Total Nucleic Acid Isolation Kit (Cat. 03038505001, Roche Diagnostic, Basel, Switzerland) following the manufacturer’s recommendations.

### RT-qPCR

The detection of RNA from ZIKV, CHIKV and DENV was carried out by RT-qPCR, using the uniplex or trioplex format.

In the case of ZIKV, primers and probes were used to amplify the coding region of protein E as previously reported [15]. Amplification of CHIKV was performed with primers and probes including positions 6856 to 6891 of the structural region of the genome [16], while for DENV, primers and probes were designed to amplify the NS5 non-structural protein [17].

The uniplex RT-qPCR assays to detect DENV were performed with the QuantiTect Probe RT-PCR Kit (Cat. 204445, Qiagen, Hilden, Germany), and the amplification of RT-PCR was performed with Superscript III Platinum One-Step reagents RT-qPCR System (Cat. 11732-088, Invitrogen, Carlsbad, CA, USA). The Applied biosystems 7500 fast platform was used in all cases. SDS software version 1.4 was used for the analysis of the results.

For the detection of RNA from ZIKV, CHIKV and DENV by trioplex RT-qPCR, the commercial VIASURE Zika, Dengue & Chikungunya Real-Time PCR Detection Kit (Cat. VS-ZDC112L, Viasure, Zaragoza, Spain) was used and implemented in the Applied biosystems 7500 fast platform and the SDS software version 2.3.

### Serology

Serum samples were processed for the determination of IgM antibodies against ZIKV and DENV by commercial qualitative methods using the commercial kits Anti-Zika Virus ELISA IgM (Cat. EI2668-9601M, Euroimmun AG, Lübeck, Germany), as well as Panbio Dengue IgM Capture ELISA (Cat. 01PE20/01PE21, Panbio Diagnostics, Korea) following the indications of the manufacturer. The results were evaluated together to discard cross-reactivity between flaviviruses (ZIKV and DENV).

The detection of IgM antibodies against Chikungunya was carried out with the reagent CHIKjj Detect IgM ELISA (Cat. CHKM-C, InBios International, Inc. USA), following the instructions of the manufacturer.

### Statistical analysis

Sample size was calculated considering published data by Cao-Lormeau et al. [9] Descriptive statistics were used for demographic and clinical data. For bivariate analysis, Pearson’s χ^2^ or Fisher exact tests were used to calculating odds ratio (OR) for paired data and 95% confidence intervals (95%CI) for the association of GBS with diagnosis by laboratory studies positive to ZIKV, DENV and CHIKV infection and symptomatology before admission. To evaluate clinical differences among groups a Mann Whitney U test was used. In all cases, a p ≤ 0.05 was considered as statistically significant.

### Ethical considerations

The study protocol was approved by the IMSS National Committee of Scientific Research (R-2016-785-026). After being provided with the necessary information and their questions clarified, all the participants who met the inclusion criteria (cases and controls) signed an informed consent form to participate in the study. A parent or guardian of any child participant provided written informed consent on their behalf.

## Results

For two years, during which the study was conducted, 1,030 cases of GBS were detected across the country. Ninety-seven cases and 184 control subjects fulfilled selection criteria and were recruited (87 cases with 2 controls and 10 cases with 1 control). Patients with GBS and control subjects were included from nine States of Mexico.

The mean age of patients with GBS and controls was 39.02 ± 19.32 years and 39.67 ± 18.49 years, respectively. Males predominated in both groups (59 and 113%, respectively), corresponding to 61% of the total participants.

### Laboratory determinations

Including all laboratory analyses (RT-qPCR in serum and urine and IgM in serum), eight ZIKV, four DENV and one CHIKV positive cases were found. ZIKV+ cases came from Veracruz and Nuevo León, and DENV+ cases came from Veracruz and Baja California. The CHIKV+ case came from Nuevo León. As for the controls, one ZIKV+ came from Veracruz), three DENV+ from Nuevo León and six CHIK+ from Veracruz (Table 1).

**Table 1.** Distribution of positive laboratory results in GBS cases (n = 97) and control subjects (n = 184) to ZIKV, DENV and CHIKV in 9 states of Mexico (July 2016 – June 2018).

| State | Cases | ZikaV + | DenV+ | ChikV+ | Controls | ZikaV + | DenV+ | ChikV+ |
| --- | --- | --- | --- | --- | --- | --- | --- | --- |
| Baja California | 10 | 0 | 1 | 0 | 19 | 0 | 0 | 0 |
| Campeche | 1 | 0 | 0 | 0 | 2 | 0 | 0 | 0 |
| Morelos | 1 | 0 | 0 | 0 | 2 | 0 | 0 | 0 |
| Chiapas | 1 | 0 | 0 | 0 | 2 | 0 | 0 | 0 |
| Nayarit | 1 | 0 | 0 | 0 | 2 | 0 | 0 | 0 |
| Nuevo León | 37 | 3 | 0 | 1 | 72 | 0 | 3 | 0 |
| Tabasco | 1 | 0 | 0 | 0 | 2 | 0 | 0 | 0 |
| Quintana Roo | 1 | 0 | 0 | 0 | 1 | 0 | 0 | 0 |
| Veracruz | 44 | 5 | 3 | 0 | 82 | 1 | 0 | 6 |
| **Total** | **97** | **8** | **4** | **1** | **184** | **1** | **3** | **6** |

An epidemic curve of GBS cases and chikungunya, dengue and Zika probable cases in the whole country during the study period is presented in Fig 1.

From the eight ZIKV+ cases, four were identified by RT-qPCR (two from serum and two from urine samples) and four were identified by IgM; of which six were DENV- and CHIKV-. Unfortunately, the remaining two samples could not be analysed due insufficient volume. All four DENV+ cases were determined by IgM; neither was positive for ZIKV nor CHIKV. The only CHIK+ case identified was negative for both ZIKV and DENV.

Using RT-qPCR, a positive ZIKV acute infection was demonstrated in four cases (two in serum and two in urine) and one control (urine) (OR, 8.04; 95% CI, 0.89-73.01; p = 0.047). In contrast, no positive DENV acute infections were found in either cases or controls. Interestingly, one positive CHIKV from the control group was identified, whereas none from the cases (Table 2).

**Table 2.**
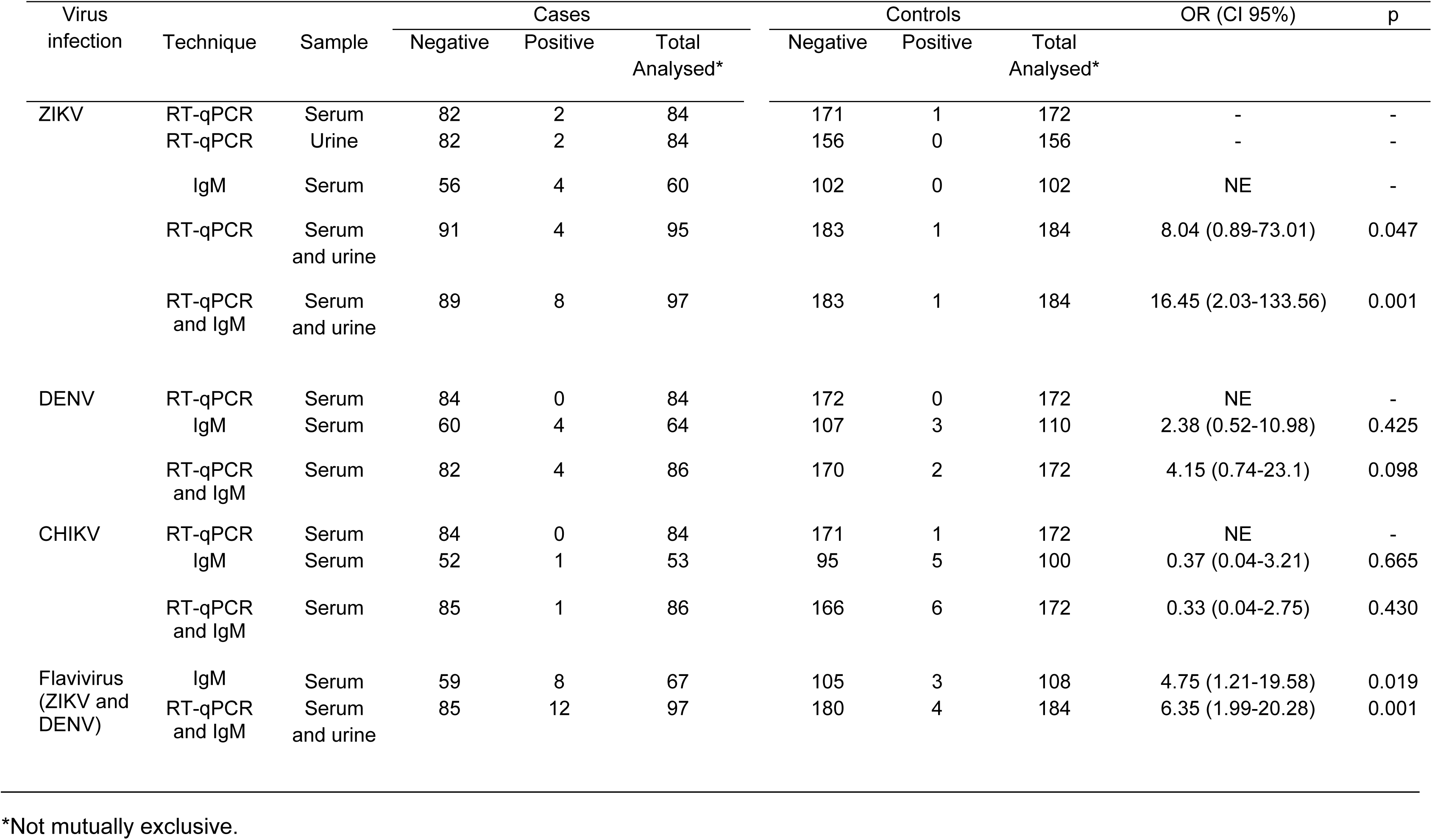
Laboratory analyses in GBS cases and controls for ZIKV, DENV or CHIKV infection according to the techniques used (RT-qPCR for serum and urine samples and IgM in serum). Mexico, July 2016-June 2018.

Evidence of recent infection by ZIKV was observed in eight GBS cases (four by RT-qPCR and four by IgM) and one control (by RT-qPCR) (OR, 16.68; 95% CI, 2.05-135.31; p = 0.001). No significant differences were observed when considering DENV+ or CHIKV+ results. However, when flaviviruses associated with GBS (ZIKV+ and DENV+) were considered toghether (12 cases = 8 ZIKV+ and 4 DENV+; and 4 controls = 1 ZIKV+ and 3 DENV+), a significant difference was observed (OR, 5.8; 95% CI, 1.8-18.8; p = 0.002). Evidence of flavivirus infection including only IgM positive results (4 ZIKV+ and 4 DENV+ cases, and 3 DENV+ controls) showed a significant difference as well (OR, 4.68; 95% CI, 1.2-18.25; p = 0.02).

### Neurologic alterations

All cases (97) were classified according to Brighton criteria (16 in level 1; 39 in level 2; and 42 in level 3). The neurological symptomatology at the nadir of all GBS cases, ZIKV+, DENV+ and ZIKV-/DENV-are shown in Table 3.

**Table 3.** Comparison of the principal signs and symptoms in the nadir of the clinical evolution of Guillain-Barré syndrome (GBS), negative, positive ZIKV and DENV, and all GBS cases. Mexico, July 2016-Jun 2018.

| Signs and symptoms | A<br>All GBS cases<br>n = 97 |  | B<br>ZIKV-/DENV-<br>n = 85 |  | C<br>ZIKV+<br>n = 8 |  | D<br>DENV+<br>n = 4 |  | p-value |  |
| --- | --- | --- | --- | --- | --- | --- | --- | --- | --- | --- |
|  | Frequency | % | Frequency | % | Frequency | % | Frequency | % | B<br>versus<br>C | B<br>versus<br>D |
| Oculomotor dysfunction | 19 | 19.6 | 14 | 16.5 | 4 | 50 | 1 | 25 | 0.02 | 0.65 |
| Facial paresis | 32 | 32 | 20 | 23.5 | 5 | 62.5 | 1 | 25 | 0.01 | 0.94 |
| Bulbar dysfunction | 21 | 21 | 13 | 18.8 | 4 | 50 | 0 | 0 | 0.04 | 0.42 |
| Upper limb weakness | 74 | 76.3 | 65 | 76.5 | 6 | 75 | 3 | 75 | 0.92 | 0.94 |
| Lower limb weakness | 89 | 90 | 74 | 87.1 | 8 | 100 | 4 | 100 | 0.28 | 0.44 |
| Generalized hypotonia | 45 | 45 | 27 | 39.1 | 7 | 87.5 | 3 | 75 | 0.01 | 0.15 |
| Dizziness | 4 | 4.1 | 2 | 2.4 | 1 | 12.5 | 1 | 25 | 0.12 | 0.01 |
| Ataxia | 21 | 21.6 | 16 | 18.8 | 2 | 25 | 3 | 75 | 0.67 | <0.01 |
| Low back pain | 3 | 3 | 2 | 2.4 | 1 | 12.5 | 0 | 0 | 0.12 | 0.75 |
| Respiratory distress | 20 | 20.6 | 16 | 18.8 | 3 | 37.5 | 1 | 25 | 0.21 | 0.76 |
| Sensory alterations | 37 | 38.1 | 32 | 37.6 | 3 | 37.5 | 2 | 50 | 0.99 | 0.62 |
| Palpitations | 2 | 2 | 2 | 2.4 | 0 | 0 | 0 | 0 | 0.66 | 0.75 |
| Low blood pressure | 2 | 2 | 1 | 1.2 | 0 | 0 | 1 | 25 | 0.75 | <0.01 |

Significant differences can be observed between ZIKV+ and ZIKV-/DENV-cases in oculomotor, facial and bulbar cranial nerve alterations and global hypotonia (p = 0.02, 0.01, 0.04 and 0.01, respectively), as well as between DENV+ and ZIKV-/DENV-cases in dizziness, ataxia and low blood pressure (p = 0.01, <0.01 and <0.01, respectively). Furthermore, a significant impairment was observed in some clinical outcomes between ZIKV+ and ZIKV-/DENV-but not with DENV+ cases: MRC at discharge (p < 0.01), GBS Disability Score at discharge (p = 0.02), and Modified Rankin Scale mainly at discharge (p = 0.02) (Table 4).

**Table 4.** Clinical characteristics and outcome of ZIKV-/DENV-, ZIKV+, DENV+ and all cases during the evolution of the Guillain-Barré syndrome in Mexico. July 2016-June 2018.

| Clinical characteristics and outcome | A<br>All GBS cases<br>n = 97 |  | B<br>ZIKV-/DENV-<br>n = 85 |  | C<br>ZIKV+<br>n = 8 |  | D<br>DENV+<br>n = 4 |  | p-value |  |
| --- | --- | --- | --- | --- | --- | --- | --- | --- | --- | --- |
|  | Mean | SD | Mean | SD | Mean | SD | Mean | SD | B<br>versus<br>C | B<br>versus<br>D |
| Latency (days) | 5.64 | 6.77 | 5.92 | 7.13 | 4.13 | 3.48 | 3.25 | 3.86 | 0.48 | 0.46 |
| Hospital stay (days) | 11.97 | 10.34 | 11.77 | 10.67 | 15.43 | 9.41 | 7.67 | 3.05 | 0.39 | 0.51 |
| MRC (admission) | 32.66 | 17.47 | 33.57 | 17.42 | 23.5 | 13.3 | 36 | 24 | 0.11 | 0.79 |
| MRC (discharge) | 42 | 17.13 | 41.51 | 16.22 | 20.67 | 14.67 | 39.33 | 26.1 | <0.01 | 0.82 |
| GBS disability score (admission) | 3.85 | 0.91 | 3.79 | 0.95 | 4.25 | 0.7 | 4.25 | 0.5 | 0.18 | 0.33 |
| GBS disability score (discharge) | 3.15 | 1.13 | 3.03 | 1.08 | 4 | 1.41 | 3.5 | 0.57 | 0.02 | 0.39 |
| Modified Rankin Scale (admission) | 3.98 | 0.96 | 3.86 | 0.97 | 4.63 | 0.51 | 4.75 | 0.5 | 0.03 | 0.07 |
| Modified Rankin Scale (discharge) | 3.25 | 1.17 | 3.13 | 1.13 | 4.13 | 1.45 | 3.5 | 0.57 | 0.02 | 0.52 |
|  | Frequency | % | Frequency | % | Frequency | % | Frequency | % |  |  |
| IV Immunoglobulin | 81 | 81 | 72 | 86.7 | 6 | 75 | 3 | 75 | 0.36 | 0.50 |
| Plasmapheresis | 2 | 2 | 2 | 3.4 | 0 | 0 | 0 | 0 | 0.71 | 0.71 |
| Mechanical ventilation | 17 | 17 | 13 | 15.7 | 3 | 42.9 | 1 | 25 | 0.07 | 0.62 |
| Nosocomial pneumonia | 2 | 2 | 2 | 3.4 | 1 | 12.5 | 0 | 0 | 0.13 | 0.54 |
| ICU management | 16 | 16.5 | 14 | 19.4 | 1 | 12.5 | 1 | 25 | 0.67 | 0.85 |
| Mortality | 4 | 4.1 | 3 | 4.9 | 1 | 12.5 | 0 | 0 | 0.46 | 0.61 |

GBS: Guillain Barré Syndrome Disability Sum Score; ICU, intensive care unit; MRC: Medical Research Council Scale for Muscle Strength Sum Score.

Intravenous immunoglobulin (81 cases) and plasmapheresis (two cases) were administered as a treatment. Mechanical ventilation was required in 17 patients (17%), of which three were ZIKV+ (42.9%) and one DENV+ (25%) (p = 0.07 and 0.62, respectively). Four deaths were reported, one for the ZIKV+ group (12.5%) and three for the ZIKV-/DENV-group (4.9%). Other clinical outcomes of GBS are shown in Table 4.

Forty electrophysiological studies were conducted, of which four were ZIKV+ cases (three of the axonal type, acute motor-sensory axonal neuropathy [AMSAN] subtype, and one, demyelinating type, acute inflammatory demyelinating polyneuropathy [AIDP] subtype); and 36 were ZIKV-/DENV-cases: 28 of the axonal type (15 AMSAN and 13 acute motor axonal neuropathy [AMAN] subtypes), and eight, demyelinating type (AIDP subtype).

In comparison with their respective controls, the cases showed significant differences in the following signs and symptoms, which were present two months before admission: rash, conjunctival hyperemia, acute diarrheal diseases (ADD), retro-ocular pain, myalgias, arthralgias, fever and headache. Other significant alterations are shown in Table 5.

**Table 5.** Symptoms prior to the development of Guillain-Barré syndrome in cases (positive or negative to ZIKV and DENV) and controls in Mexico. July 2016-June 2018.

| Symptoms | Controls<br>n = 184 |  | GBS cases<br>n = 97 |  | OR (CI 95%) | p-value | ZIKV-/DENV-<br>n = 85 |  | ZIKV+<br>n = 8 |  | DENV+<br>n = 4 |  |
| --- | --- | --- | --- | --- | --- | --- | --- | --- | --- | --- | --- | --- |
|  | Frequency | % | Frequency | % |  |  | Frequency | % | Frequency | % | Frequency | % |
| Rash | 2 | 1 | 22 | 22.7 | 26.69 (6.12-116.36) | p<0.01 | 18 | 21 | 3 | 37.5 | 1 | 25 |
| Conjunctival hyperemia | 2 | 1 | 18 | 18.6 | 20.73 (4.69-91.49) | p<0.01 | 15 | 18 | 3 | 37.5 | 0 | 0 |
| ADD | 2 | 1 | 13 | 13.4 | 16.34 (3.59-74.36) | p<0.01 | 13 | 15 | 0 | 0 | 0 | 0 |
| Retro-ocular pain | 2 | 1 | 11 | 11 | 11.64 (2.52-53.66) | p<0.01 | 8 | 9.4 | 3 | 37.5 | 0 | 0 |
| Myalgias | 12 | 6.5 | 41 | 42 | 10.49 (5.15-21.35) | p<0.01 | 34 | 40 | 4 | 50 | 3 | 75 |
| Diarrhea | 10 | 5.4 | 35 | 36 | 9.82 (4.59-21.00) | p<0.01 | 32 | 38 | 2 | 25 | 1 | 25 |
| Arthralgias | 12 | 6.5 | 39 | 40 | 9.63 (4.72-19.64) | p<0.01 | 34 | 40 | 3 | 37.5 | 2 | 50 |
| Malaise | 16 | 8.7 | 46 | 47.4 | 9.47 (4.94-18.13) | p<0.01 | 40 | 47 | 4 | 50 | 2 | 50 |
| Pruritus | 3 | 1.6 | 12 | 12 | 8.51 (2.34-30.97) | p<0.01 | 9 | 10 | 3 | 37.5 | 0 | 0 |
| Headache | 21 | 11.4 | 49 | 50.5 | 7.92 (4.31-14.49) | p<0.01 | 42 | 49 | 4 | 50 | 3 | 75 |
| Fever | 18 | 9.8 | 44 | 45 | 7.65 (4.07-14.36) | p<0.01 | 36 | 42 | 6 | 75 | 3 | 75 |
| Sore throat | 6 | 3.3 | 14 | 14.4 | 5.00 (1.85-13.48) | p<0.01 | 10 | 12 | 3 | 37.5 | 1 | 25 |
| ARI | 8 | 4.3 | 13 | 13.4 | 3.74 (1.48-9.43) | p<0.01 | 12 | 14 | 1 | 12.5 | 0 | 0 |
| Cough | 11 | 6 | 16 | 16.5 | 3.10 (1.38-6.99) | p<0.01 | 15 | 18 | 1 | 12.5 | 0 | 0 |
| Vomiting | 8 | 4.3 | 11 | 11 | 2.81 (1.09-7.25) | p 0.02 | 10 | 12 | 0 | 0 | 1 | 25 |
| Nausea | 9 | 5 | 12 | 12.4 | 2.74 (1.11-6.76) | p 0.02 | 11 | 13 | 0 | 0 | 1 | 25 |
| Abdominal pain | 17 | 9.2 | 21 | 21.6 | 2.71 (1.35-5.43) | p<0.01 | 18 | 21 | 1 | 12.5 | 2 | 50 |
| Expectoration | 4 | 2 | 5 | 5 | 2.44 (0.64-9.32) | p 0.17 | 4 | 5 | 1 | 12.5 | 0 | 0 |
| Nasal congestion | 8 | 4 | 9 | 9.3 | 2.25 (0.83-6.03) | p 0.10 | 9 | 10 | 0 | 0 | 0 | 0 |
| Rhinorrhea | 10 | 5.4 | 11 | 11 | 2.22 (0.91-5.44) | p 0.07 | 11 | 13 | 0 | 0 | 0 | 0 |
| Constipation | 4 | 2.2 | 3 | 3 | 1.43 (0.35-6.55) | p 0.63 | 2 | 2 | 1 | 12.5 | 0 | 0 |
| Coryza | 2 | 1 | 1 | 1 | 0.94 (0.08-10.58) | p 0.96 | 1 | 1 | 0 | 0 | 0 | 0 |
| Lymphadenopathy | 2 | 1 | 1 | 1 | 0.94 (0.08-10.58) | p 0.96 | 0 | 0 | 1 | 12.5 | 0 | 0 |
| Chest pain | 4 | 2 | 2 | 2 | 0.94 (0.17-5.26) | p 0.95 | 2 | 2 | 0 |  | 0 | 0 |
| Pelvic pain | 5 | 2.7 | 2 | 2 | 0.75 (0.14-3.95) | p 0.73 | 2 | 2 | 0 | 0 | 0 | 0 |
| Abdominal distension | 12 | 6.5 | 3 | 3 | 0.45 (0.12-1.66) | p 0.22 | 2 | 2 | 1 | 12.5 | 0 | 0 |

ADD: acute diarrheal diseases; ARI: acute respiratory infections; CI 95%: confidence interval 95%; OR: odds ratio.

## Discussion

The present study results show an evident association between GBS and ZIKV, as well as between GBS and DENV, both flaviviruses, supported by laboratory analysis. Moreover, greater severity of GBS is associated with ZIKV and co-infections were not demonstrated among these three viruses.

Several reports indicate the probability of a causal association between ZIKV and GBS due to the increase in the number of cases of GBS during ZIKV outbreaks [18-20]. Some describe a greater severity of GBS symptoms [12,13,21]. However, only seven reports with an appropriate design to show a positive or negative association were found [9,21-26].

In the present work, viral RNA showing ZIKV acute infection was detected (OR, 8.04; 95% CI, 0.89-73.01; p = 0.047). Although the lower endpoint of the confidence interval is < 1, the preponderance of evidence suggests an association between ZIKV infection and GBS. At present, only one report has demonstrated an acute ZIKV infection associated with GBS by RT-PCR, in Puerto Rico, although the biological sample (serum, urine, saliva) was not specific [22].

When considering the evidence of ZIKV recent infection by RT-PCR (serum or urine samples) or IgM anti-ZIKV (serum), our findings demonstrated an association with GBS, similar to Dirlikov et al., in Puerto Rico [22], and Simon et al., in New Caledonia [25]. Moreover, previous infection by ZIKV was also considered through the combination of positive results with IgM and IgG anti ZIKV, along with the neutralising response against ZIKV, described by Cao-Lomeau et al. in French Polynesia, demonstrating the association between GBS and ZIKV for the first time [9].

Concerning GBS secondary to acute DENV infection identified by RT-PCR, most publications indicate an apparent association [5,27,28]. Unfortunately, a causal association has not been convincingly demonstrated. In the present research, this association could not be demonstrated either. However, a significant association between GBS and flavivirus (ZIKV and DENV) was demonstrated, either by combining RT-PCR and IgM results or by using only IgM results. These findings agree with those previously reported by Salinas et al. [23], Stycinski et al. [24] and Simon et al. [25] in other populations (Colombia, Brazil, and New Caledonia, respectively). Although no conclusive role of DENV in the triggering of GBS has been demonstrated, an association between GBS and flavivirus (ZIKV and DENV) seems to be established.

No association between GBS and CHIKV could be demonstrated, probably due to the small sample size or a less CHIKV infection incidence in Mexico (Figure 1). Finally, it was not possible to categorically discard any co-infection cases, because although 11 of 13 of positive ZIKV, DENV or CHIKV patients were negative to the other two viruses, two of ZIKV+ patients showed no DENV or CHIKV results. However, it is proposed that no co-infection cases were present in this samples, which disagrees with other results described before [6,7].

**Fig 1.**
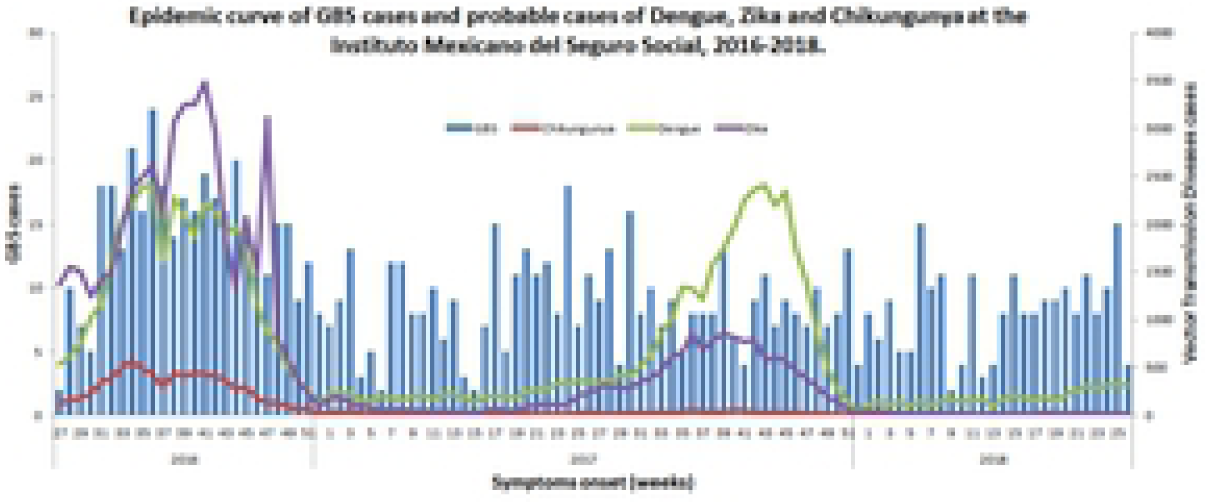
Epidemic curve of GBS cases and probable cases of dengue, Zika and chikungunya at Instituto Mexicano del Seguro Social (IMSS). Mexico July 2016-June 2018. Source: National System of Epidemiological Surveillance of Acute Flaccid Paralysis and Vector Transmission Diseases, and Morbidity Yearbook; Ministry of Health, Mexico. http://187.191.75.115/anuario/2016/incidencia/incidencia_casos_nuevos_enfermedad_grupo_edad.pdf http://187.191.75.115/anuario/2017/incidencia/incidencia_casos_nuevos_enfermedad_grupo_edad.pdfhttp://www.paho.org/data/index.php/es/temas/indicadores-dengue/dengue-nacional/240-dengue-incidencia.html?start=3 https://www.paho.org/hq/index.php?option=com_docman&view=download&category_slug=2016-8380&alias=37868-numero-casos-reportados-fiebre-chikungunya-americas-2016-868&Itemid=270&lang=es http://www.paho.org/data/index.php/es/temas/indicadores-dengue/dengue-nacional/240-dengue-incidencia.html?start=3

### Neurological behaviour

GBS associated with ZIKV is often related to dysautonomia, paralysis of cranial nerves and more rapid onset of signs and symptoms [20,21]. However, in the present work, no dysautonomia was observed, but the paralysis of cranial nerves (oculomotor, facial and bulbar nerves paresis) and generalised hypotonia were the most significant alterations observed. The need for ventilator support varies from to 28% according to other studies [14,29]. In contrast, it was more common in the present work (42.9%). Although the difference was not statistically significant, it is consistent with previous reports [21]. Finally, although the mortality of GBS cases in the present work (4.1%) is within the values reported in other studies in the world and Mexico [10,11,14,29,30], it was higher for the cases associated with ZIKV (12.5%). A higher percentage of patients with muscular weakness, impairment of functional status and incapacity, as well as higher percentage of mechanical ventilation, and relatively higher mortality shown in the present study indicate that GBS cases associated with ZIKV may be more severe than those seen with other antecedent etiologies of GBS [31,32].

Considering the electrophysiological results of the present cases, AMSAN subtype was more frequently observed, which agrees with what has been reported for the Asia/Asia Pacific (French Polynesia) [9] and the Americas (Cúcuta, Colombia) [20] regions, where the axonal type also predominated but AMAN subtype. On the contrary, these results differ from other regions of Asia, including Bangladesh and the Americas (Cúcuta and Barranquilla, in Colombia) as well, in which the demyelinating type AIDP subtype was the most frequent [21,23,26].

In this work, symptoms prior to the development of GBS related to the symptoms of ZIKV suspected disease, such as rash, fever, conjunctivitis and arthralgias showed significant differences with respect to controls. Other authors have highlighted these symptoms as symptoms of suspicion [23,24]. In 2016, Cao-Lormeau et al. described these symptoms (rash, arthralgia and fever); however, they did not analyse the risk [9]. Some recent studies have reported this association: Salinas et al. (2017) reported rash, fever, arthralgia and myalgia [23] and Styczynski et al. (2017) reported rash, headache, fever, arthralgias and myalgias [24]. When considering the symptoms reported in these studies and including our findings, the most significant symptoms are rash, arthralgia, fever, myalgia, conjunctivitis and headache.

The symptoms of DENV suspected disease, such as fever, headache, arthralgias, myalgias, rash, nausea and vomiting have been described previously [5]. In the present work, they were also significantly different when compared with controls. All these results support the association between GBS and ZIKV and DENV infection.

Considering that ZIKV, DENV, and CHIKV produce large epidemics worldwide [1] and these three arboviruses coexist in Mexico since 2015 [33], this study confirms their presence in both GBS cases and controls. However, only ZIKV or DENV but no CHIKV were associated with GBS. Similar to previous studies, the present report also demonstrated that GBS cases associated with ZIKV showed a more severe clinical picture [12, 13)], which should be considered for the treatment of these patients in countries where previous ZIKV infection could originate this neurological disease. Although co-infections between these arboviruses have been described previously [6,7], cases in which two or three of these viruses were simultaneously present were not demonstrated in this study. Finally, while the incidence of GBS in Mexico was not aimed on the present study, unpublished data of the Instituto Mexicano del Seguro Social (the Mexican Institute of Social Security) shows that the average of GBS cases/year increased 23% during 2016-2018 in comparison with 2013-2015, which could be associated with an increased number of ZIKV cases, given that DENV and CHIKV had been previously identified in Mexico.

Some limitations of the study were the small sample size obtained for an epidemiological study, although the study was based on an Institutional Epidemiological Surveillance of Communicable Diseases system. Researches decided voluntarily to participate in the study. The participation was not mandatory. Finally, in some cases, it was not possible to demonstrate ZIKV infection in patients with GBS due to a delay in medical attention.

In conclusion, the present study demonstrates that acute ZIKV infection is associated with GBS as well as the laboratory evidence of the association of GBS with both a recent ZIKV infection and a recent flavivirus infection (ZIKV or DENV). On the one hand, no association between GBS and CHIKV infection was found neither co-infections were demonstrated among these three viruses. On the other hand, GBS cases associated with ZIKV showed a more severe clinical picture (more significant impairment of functional status and incapacity, a higher percentage of mechanical ventilation and mortality). GBS associated with DENV cases seemed to show more dizziness, ataxia and low blood pressure. Finally, the symptoms of ZIKV or DENV suspected disease were observed before the development of GBS, which is consistent with other studies published previously and support the association between GBS and ZIKV and DENV infection.

## Acknowledgments

The authors would like to thank to Dr. James Sejvar and Dr. Rosalia Lira Carmona because their importat commentaries to the manuscript. We also would like to thank to Anel Cantera Salinas, Osvaldo Jiménez Pacheco, Vidaris Toledo Arrazola and Mario Villegas Rivera, who helped us to elaborate the clinical data base.

## Supporting Information Legends

S1 Appendix: STROBE Checklist

## References

1. Weaver SC, Reisen WK. Present and future arboviral threats. Antiviral Res. 2010;85(2): 328. doi: 10.1016/j.antiviral.2009.10.008

2. Patterson J, Sammon M, Garg M. Dengue, Zika and chikungunya: emerging arboviruses in the New World. West J Emerg Med. 2016;17(6):671–9. doi: 10.5811/westjem.2016.9.30904

3. Broutet N, Krauer F, Riesen M, Khalakdina A, Almiron M, Aldighieri S, et al. Zika virus as a cause of neurologic disorders. N Engl J Med. 2016;374(16):1506–9. doi: 10.1056/NEJMp1602708

4. Mehta R, Gerardin P, de Brito CAA, Soares CN, Ferreira MLB, Solomon T. The neurological complications of chikungunya virus: a systematic review. Rev Med Virol. 2018;28(3):e1978. doi: 10.1002/rmv.1978

5. Verma R, Sahu R, Holla V. Neurological manifestations of dengue infection: a review. J Neurol Sci. 2014;346(1-2):26–34. doi: 10.1016/j.jns.2014.08.044

6. Acevedo N, Waggoner J, Rodriguez M, Rivera L, Landivar J, Pinsky B, et al. Zika virus, chikungunya virus, and dengue virus in cerebrospinal fluid from adults with neurological manifestations, Guayaquil, Ecuador. Front Microbiol. 2017;8:42. doi: 10.3389/fmicb.2017.00042

7. Azevedo MB, Coutinho MSC, Silva MAD, Arduini DB, Lima JDV, Monteiro R, et al. Neurologic manifestations in emerging arboviral diseases in Rio de Janeiro City, Brazil, 2015-2016. Rev Soc Bras Med Trop. 2018;51(3):347–51. doi: 10.1590/0037-8682-0327-2017

8. Willison HJ, Jacobs BC, van Doorn PA. Guillain-Barré syndrome. Lancet 2016;388(10045):717–27. doi: 10.1016/S0140-6736(16)00339-1

9. Cao-Lormeau VM, Blake A, Mons S, Lastère S, Roche C, Vanhomwegen J, et al. Guillain–Barré syndrome outbreak associated with Zika virus infection in French Polynesia: a case–control study. Lancet 2016;387(10027):1531–9. doi: 10.1016/S0140-6736(16)00562-6

10. Souayah N, Nasar A, Suri MFK, Qureshi AI. National trends in hospital outcomes among patients with Guillain-Barré syndrome requiring mechanical ventilation. J Clin Neuromuscul Dis. 2008;10(1):24–8. doi: 10.1097/CND.0b013e3181850691

11. Dominguez-Moreno R, Tolosa Tort P, Patiño-Tamez A, Quintero-Bauman A, Collado-Frias D, Miranda-Rodríguez MG, et al. Mortality associated with a diagnosis of Guillain-Barre syndrome in adults of Mexican health institutions. Rev Neurol. 2014;58(1):4–10. doi: https://doi.org/10.33588/rn.5801.2013370

12. Paul LM, Carlin ER, Jenkins MM, Tan AL, Barcellona CM, Nicholson CO, et al. Dengue virus antibodies enhance Zika virus infection. Clin Transl Immunology 2016;5(12):e117. doi:10.1038/cti.2016.72

13. Castanha PMS, Nascimento EJM, Braga C, Cordeiro MT, de Carvalho OV, de Mendonça LR, et al. Dengue virus-specific antibodies enhance Brazilian Zika virus infection. J Infect Dis. 2017;215:781–5. doi: 10.1093/infdis/jiw638

14. Fokke C, van den Berg B, Drenthen J, Walgaard C, van Doorn PA, Jacobs BC. Diagnosis of Guillain-Barré syndrome and validation of Brighton criteria. Brain 2014;137:33–43. doi: 10.1093/brain/awt285

15. Lanciotti RS, Kosoy OL, Laven JJ, Velez JO, Lambert AJ, Johnson AJ, et al. Genetic and serologic properties of Zika virus associated with an epidemic, Yap State, Micronesia, 2007. Emerg Infect Dis. 2008;14(8):1232–9. doi: 10.3201/eid1408.080287

16. Lanciotti RS, Kosoy OL, Laven JJ, Panella AJ, Velez JO, Lambert AJ, et al. Chikungunya virus in US travelers returning from India, 2006. Emerg Infect Dis. 2007;13(5):764–7. doi: 10.3201/eid1305.070015

17. Chien LJ, Liao TL, Shu PY, Huang JH, Gubler DJ, Chang GJ. Development of real-time reverse transcriptase PCR assays to detect and serotype dengue viruses. J Clin Microbiol. 2006;44(4):1295–304. doi: 10.1128/JCM.44.4.1295-1304.2006

18. Parra B, Lizarazo J, Jiménez-Arango JA, Zea-Vera AF, González-Manrique G, Vargas J, et al. Guillain–Barré syndrome associated with Zika virus infection in Colombia. N Engl J Med. 2016;375(16):1513–23. doi: 10.1056/NEJMoa1605564

19. Barbi L, Coelho AVC, Alencar LCA, Crovella S. Prevalence of Guillain-Barré syndrome among Zika virus infected cases: a systematic review and meta-analysis. Braz J Infect Dis. 2018;22(2):137–41. doi: 10.1016/j.bjid.2018.02.005

20. Arias A, Torres-Tobar L, Hernandez G, Paipilla D, Palacios E, Torres Y, et al. Guillain-Barre syndrome in patients with a recent history of Zika in Cúcuta, Colombia: a descriptive case series of 19 patients from December 2015 to March 2016. J Crit Care. 2017;37:19–23. doi:10.1016/j.jcrc.2016.08.016

21. Anaya JM, Rodriguez Y, Monsalve DM, Vega D, Ojeda E, Gonzalez-Bravo D, et al. A comprehensive analysis and immunobiology of autoimmune neurological syndromes during the Zika virus outbreak in Cúcuta, Colombia. J Autoimmun. 2017;77:123–38. doi:10.1016/j.jaut.2016.12.007

22. Dirlikov E, Medina NA, Major CG, Munoz-Jordan JL, Luciano CA, Rivera-Garcia B, et al. Acute Zika virus Infection as a risk factor for Guillain-Barre syndrome in Puerto Rico. JAMA. 2017;318(15):1498–500. doi:10.1001/jama.2017.11483

23. Salinas JL, Walteros DM, Styczynski A, Garzon F, Quijada H, Bravo E, et al. Zika virus disease-associated Guillain-Barre syndrome-Barranquilla, Colombia 2015-2016. J Neurol Sci. 2017;381:272–7. doi:10.1016/j.jns.2017.09.001

24. Styczynski AR, Malta J, Krow-Lucal ER, Percio J, Nobrega ME, Vargas A, et al. Increased rates of Guillain-Barre syndrome associated with Zika virus outbreak in the Salvador metropolitan area, Brazil. PLoS Negl Trop Dis. 2017;11(8):e0005869. doi:10.1371/journal.pntd.0005869

25. Simon O, Acket B, Forfait C, Girault D, Gourinat AC, Millon P, et al. Zika virus outbreak in New Caledonia and Guillain-Barre syndrome: a case-control study. J Neurovirol. 2018; 10.1007/s13365-018-0621-9. doi:10.1007/s13365-018-0621-9

26. GeurtsvanKessel CH, Islam Z, Islam MB, Kamga S, Papri N, van de Vijver D, et al. Zika virus and Guillain-Barre syndrome in Bangladesh. Ann Clin Transl Neurol. 2018;5(5):606–15. doi:10.1002/acn3.556

27. Simon O, Billot S, Guyon D, Daures M, Descloux E, Gourinat AC, et al. Early Guillain–Barre syndrome associated with acute dengue fever. J Clin Virol 2016;77:29–31. doi: 10.1016/j.jcv.2016.01.016

28. Soares CN, Cabral-Castro M, Oliveira C, Faria LC, Perarlta JM, Freitas MR, et al. Oligosymptomatic dengue infection: a potential cause of Guillain Barre syndrome. Arq Neuropsiquiatr. 2008;66(2A):234–7. doi: 10.1590/s0004-282x2008000200018

29. Alshenkhlee A, Hussain Z, Sultan B, Katirji B. Guillain-Barré síndrome: incidence and mortality rates in US hospitals. Neurology 2008;70(18):1608–13. doi: 10.1212/01.wnl.0000310983.38724.d4

30. Ancona P, Bailey M, Bellomo R. Characteristics, incidence and outcome of patients admitted to intensive care unit with Guillain-Barre syndrome in Australia and New Zealand. J Crit Care 2018;45:58–64. doi: 10.1016/j.jcrc.2018.01.016

31. Sebastián UU, Ricardo AVA, Alvarez BC, Cubides A, Luna AF, Arroyo-Parejo M, et al. Zika virus-induced neurological critical illness in Latin America: Severe Guillain-Barre Syndrome and encephalitis. J Crit Care 2017;42:275–81. doi: 10.1016/j.jcrc.2017.07.038.

32. Uncini A, Gonzáñez-Bravo DC, Acosta-Ampudia YY, Ojeda EC, Rodríguez Y, Monsalve DM, et al. Clinical and nerve conduction features in Guillain-Barré syndrome associated with Zika virus infection in Cúcita, Colombia. Eur J Neurol 2018;25(4):644–50. doi: 10.1111/ene.13552.

